# Interplay between axonal Wnt5-Vang and dendritic Wnt5-Drl/Ryk signaling controls glomerular patterning in the *Drosophila* antennal lobe

**DOI:** 10.1101/813246

**Authors:** Huey Hing, Noah Reger, Jennifer Snyder, Lee G. Fradkin

## Abstract

Despite the importance of dendritic targeting in neural circuit assembly, the mechanisms by which it is controlled still remain incompletely understood. We previously showed that in the developing *Drosophila* antennal lobe, the Wnt5 protein forms a gradient that directs the ∼45° rotation of a cluster of projection neuron (PN) dendrites, including the adjacent DA1 and VA1d dendrites. We report here that the Van Gogh (Vang) transmembrane planar cell polarity (PCP) protein is required for the rotation of the DA1/VA1d dendritic pair. Cell type-specific rescue and mosaic analyses showed that Vang functions in the olfactory receptor neurons (ORNs), suggesting a codependence of ORN axonal and PN dendritic targeting. Loss of Vang suppressed the repulsion of the VA1d dendrites by Wnt5, indicating that Wnt5 signals through Vang to direct the rotation of the DA1 and VA1d glomeruli. We observed that the Derailed (Drl)/Ryk atypical receptor tyrosine kinase is also required for the rotation of the DA1/VA1d dendritic pair. Antibody staining showed that Drl/Ryk is much more highly expressed by the DA1 dendrites than the adjacent VA1d dendrites. Mosaic and epistatic analyses showed that Drl/Ryk specifically functions in the DA1 dendrites in which it antagonizes the Wnt5-Vang repulsion and mediates the migration of the DA1 glomerulus towards Wnt5. Thus, the nascent DA1 and VA1d glomeruli appear to exhibit Drl/Ryk-dependent biphasic responses to Wnt5. Our work shows that the final patterning of the fly olfactory map is the result of an interplay between ORN axons and PN dendrites, wherein converging pre- and postsynaptic processes contribute key Wnt5 signaling components, allowing Wnt5 to orient the rotation of nascent synapses through a PCP mechanism.

## Introduction

The prevailing view of neural circuit assembly is that axons and dendrites are separately guided by molecular gradients to their respective positions whereupon they form synapses with each other (1–4). However, careful observation of developing neural circuits reveals that the process may be more complex. For example, in the developing retina outer plexiform layer (OPL) the axon terminals of rods and cones, and dendrites of their respective postsynaptic cells, the rod and cone bipolar cells, are initially intermingled in the nascent OPL (5). Even as the rod and cone axons are connecting with their target dendrites, the terminals are segregating into rod- and cone-specific sub-laminae, suggesting that the processes of targeting and synaptic partner matching may be coordinated. Whether the two processes are functionally linked and what mechanisms might be involved are unknown.

The stereotyped neural circuit of the *Drosophila* olfactory map offers a unique opportunity to unravel the mechanisms of neural circuit development. Dendrites of 50 classes of uniglomerular projection neurons (PNs) form synapses with the axons of 50 classes of olfactory receptor neurons (ORNs) in the antennal lobe (AL) in unique glomeruli (6, 7). This precise glomerular map is thought to be established during the pupal stage by the targeting of PN dendrites (8–10). We previously reported that during the establishment of the fly olfactory map, two adjacent dendritic arbors located at the dorsolateral region of the AL, the DA1 and VA1d dendrites (hereafter referred to as the DA1/VA1d dendritic pair), undergo rotational migration of ∼45° around each other to attain their final adult positions (11). This rearrangement (in the lateral/90° → dorsal/0° → medial/270° → ventral/180° direction) occurs between 16 and 30 hour After Puparium Formation (hAPF), a period of major ORN axon ingrowth to the AL (8, 12, 13). We showed that a Wnt5 signal guides this rotation by repelling the dendrites (11). Wnt5 is expressed by a set of AL-extrinsic cells and forms a dorsolateral-high to ventromedial-low (DL>VM) gradient in the AL neuropil which provides a directional cue to align the dendritic pattern relative to the axes of the brain. We also showed that the Derailed (Drl)/Ryk kinase-dead receptor tyrosine kinase, a Wnt5 receptor (14–17), is differentially expressed by the PN dendrites, thus providing cell-intrinsic information for their targeting in the Wnt5 gradient. Interestingly, *drl* opposes *Wnt5* repulsive signaling so that dendrites expressing high levels of *drl* terminate in regions of high *Wnt5* concentration and *vice versa*. To further unravel the mechanisms of PN dendritic targeting, we have screened for more mutations that disrupt the rotation of the DA1/VA1d dendritic pair.

Here we report that mutations in the *Van Gogh* (*Vang*) gene disrupted the rotation of the DA1/VA1d dendritic pair, thus mimicking the *Wnt5* mutant phenotype. *Vang* encodes a four-pass transmembrane protein (18, 19) of the core Planar Cell Polarity (PCP) group, an evolutionarily conserved signaling module that imparts polarity to cells (20, 21). The loss of *Vang* suppressed the repulsion of the VA1d dendrites by *Wnt5*, indicating that *Vang* is a downstream component of *Wnt5* signaling. Surprisingly, *Vang* acts in the ORNs, which suggests an obligatory codependence of ORN axon and PN dendritic migration. We also show that the *drl* gene is selectively expressed in the DA1 dendrites where it antagonizes *Vang* and appears to convert *Wnt5* repulsion of the DA1 glomerulus into attraction. The opposing responses of the DA1 and VA1d glomeruli likely create the forces by which *Wnt5* directs the rotation of the glomerular pair. Our work shows that converging pre- and postsynaptic processes contribute key signaling components of the *Wnt5* pathway, allowing the processes to be co-guided by the *Wnt5* signal.

## Results

### *Vang* promotes the rotation of DA1/VA1d dendrites

We have shown that during wild-type development the adjacent DA1 and VA1d dendrites rotate around each other, such that DA1 moves from its original position lateral to VA1d at 18 hAPF to its final position dorsolateral to VA1d in the adult, an ∼45° rotation (11). We also showed that this rotation requires the *Wnt5* gene, for in the null *Wnt5^400^* mutant the rotation is abolished, resulting in an adult DA1/VA1d angle of 76.03° ± 3.6° (N=29, vs 29.32° ± 2.5°, N=22 in the *Wnt5^400^/+* heterozygous control, Student’s *t* test p<0.0001) (See Methods for quantification) (Fig 1A-C). The DA1 and VA1d pair of dendrites were visualized by expressing *UAS-mCD8::GFP* under the control of *Mz19-Gal4*, which specifically labels the DA1, VA1d and DC3 dendrites (22, 23). To elucidate the molecular mechanisms by which *Wnt5* controls the rotation of the PN dendrites, we screened a panel of signal transduction mutants for similar defects in DA1/VA1d rotation. We found that animal homozygous for *Vang* mutations exhibit a DA1/VA1d phenotype that mimicked that of the *Wnt5^400^* mutant. For example, in the null *Vang^6^* allele, the DA1/VA1d angle was 54.72° ± 2.8° (N=24, vs 27.78° ± 4.6°, N=18 in the *Vang^6^/+* heterozygous control, *t*-test p<0.0001) (Fig 1D-F) suggesting that *Vang* might function in the *Wnt5* pathway. Since the *Vang^6^* allele, which encodes a truncated 128 amino acid product (19), displayed a highly penetrant phenotype, we examined this allele further. We therefore examined the positioning of glomeruli in different regions of the *Vang^6^* AL by expressing *UAS-mCD8::GFP* under the control of various *Or-Gal4* drivers (6) (Fig 1G-O, see Methods for quantification). We observed that glomeruli in the lateral region of the AL, such as the VA1lm glomerulus (Fig 1G-I), showed the greatest displacement compared with glomeruli in other regions, suggesting that *Vang* primarily controls neurite targeting in the lateral AL. Since the Wnt5 protein is highly concentrated at the dorsolateral region of the AL (11), the *Vang* mutant defects are consistent with *Vang* playing a role in *Wnt5* signaling. We hypothesized that *Vang* mediates *Wnt5* signaling in the control of the DA1/VA1d dendritic rotation. Student’s *t* tests were used to compare the data of the mutants with those of their respective controls. Scale bars: 10 µm.

**Fig 1.**
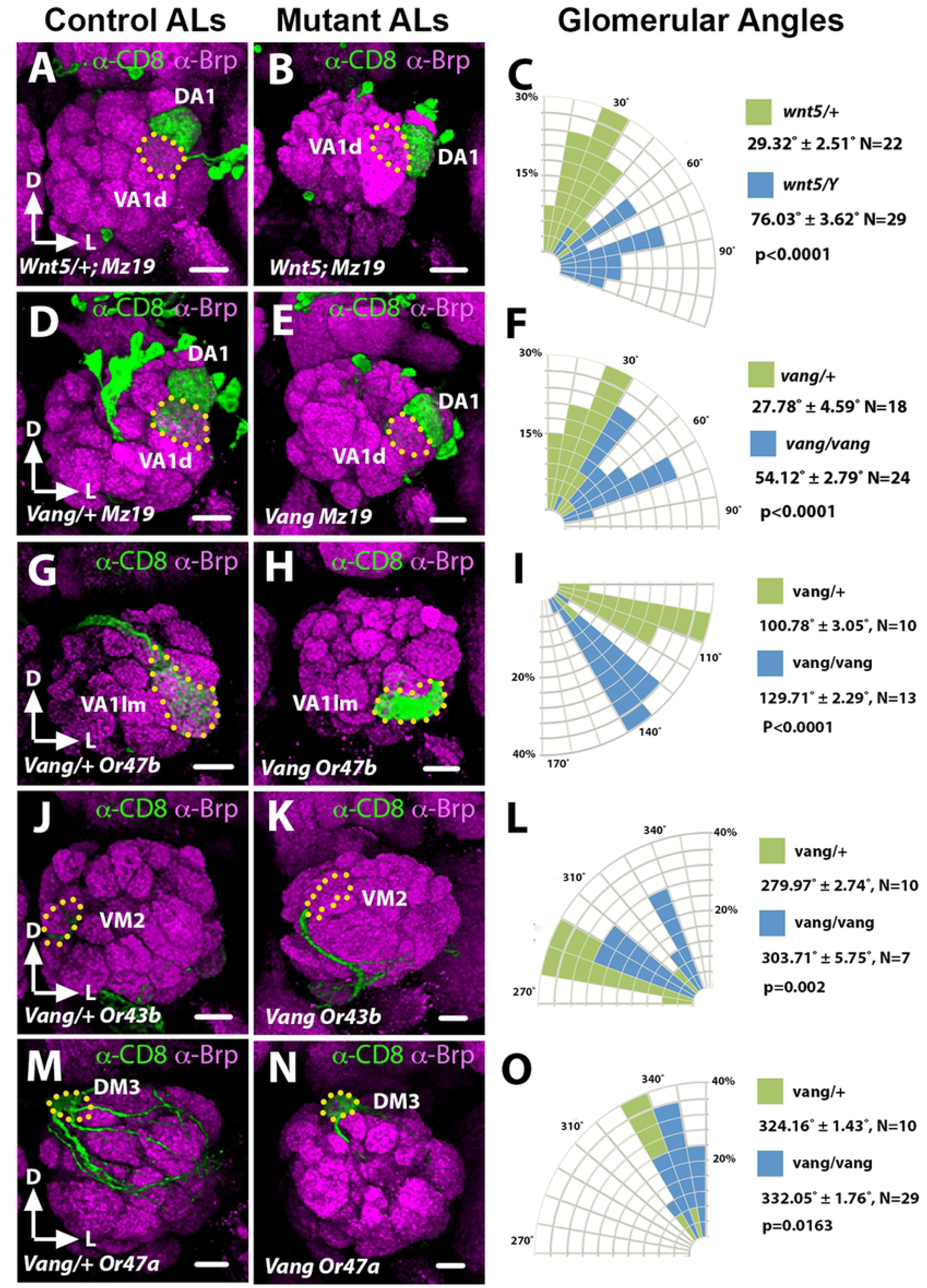
The *Vang^6^* mutant AL defects mimic that of the *Wnt5^400^* mutant. Frontal views of the left ALs are shown (dorsal up and lateral to the right) in this and following figures. (**A-F**) Adult ALs from animals expressing *UAS-mCD8::GFP* under the control of *Mz19-Gal4* stained with antibodies against Bruchpilot (Brp, Magenta) to highlight the AL neuropil and CD8 (green) to highlight the DA1 and VA1d PN dendritic arbors in the *Wnt5^400^/+* (A) and *Vang^6^/+* (D) controls and *Wnt5^400^* (B) and *Vang^6^* (E) mutants. (C, F) Quantification of the DA1/VA1d angles in the *Wnt5^400^* and *Vang^6^* mutants respectively. The DA1 dendrites are located dorsal to the VA1d dendrites in the controls but lateral to the VA1d dendrites in the *Wnt5^400^* and *Vang^6^* mutants. (**G-O**) Left adult ALs from animals expressing *UAS-mCD8::GFP* under the control of *Or47b-Gal4* (G, H), *Or43b-Gal4* (J, K) and *Or47a-Gal4* (M, N) to label lateral, medial and dorsal glomeruli respectively, in the *Vang^6^/+* (G, J, M) controls and *Vang^6^* mutants (H, K, N). (**I, L, O**) Quantification of the positions of the *Or47b*, *Or43b* and *Or47a* glomeruli respectively in the control vs *Vang^6^* animals. The glomeruli appeared to be displaced in the clockwise direction in the *Vang^6^* mutant, with VA1lm showing the greatest displacement.

To obtain further evidence for *Vang*’s role in regulating the rotation of the DA1/VA1d dendritic pair, we stained ALs during a time of active glomerular rotation (24 hAPF) (11) with an antibody directed against the N-terminal 143 amino acids of Vang (24). We observed that the Vang staining has a punctate appearance and is highly concentrated in the dorsolateral region of the AL between 0-9 µm from the AL anterior surface (Fig 2A). Co-labeling of the DA1/VA1d dendrites with the *Mz19-Gal4* driver showed that they reside between ∼3-6 µm in this high Vang expression domain. Vang staining in the neuropil begins to decline at 10 µm but strongly highlighted the nerve fiber layer (arrow) and the antennal nerve at 8-12 µm (arrow and arrowheads in Fig 2A and B), as well as the antennal commissure at 22 µm depth. The antibody stained the *Vang^6^* mutant ALs (Fig 2C), likely because the *Vang^6^* allele encodes a truncated protein. Nonetheless, the strong reduction in staining intensity compared with wild-type ALs attested to the antibody’s specificity. We concluded that Vang is expressed in the AL during the period of active AL neuropil rotation, where it colocalized with the DA1 and VA1d dendrites. The Vang expression pattern is consistent with the hypothesis that *Vang* mediates *Wnt5* control of the DA1/VA1d dendritic rotation.

**Fig 2.**
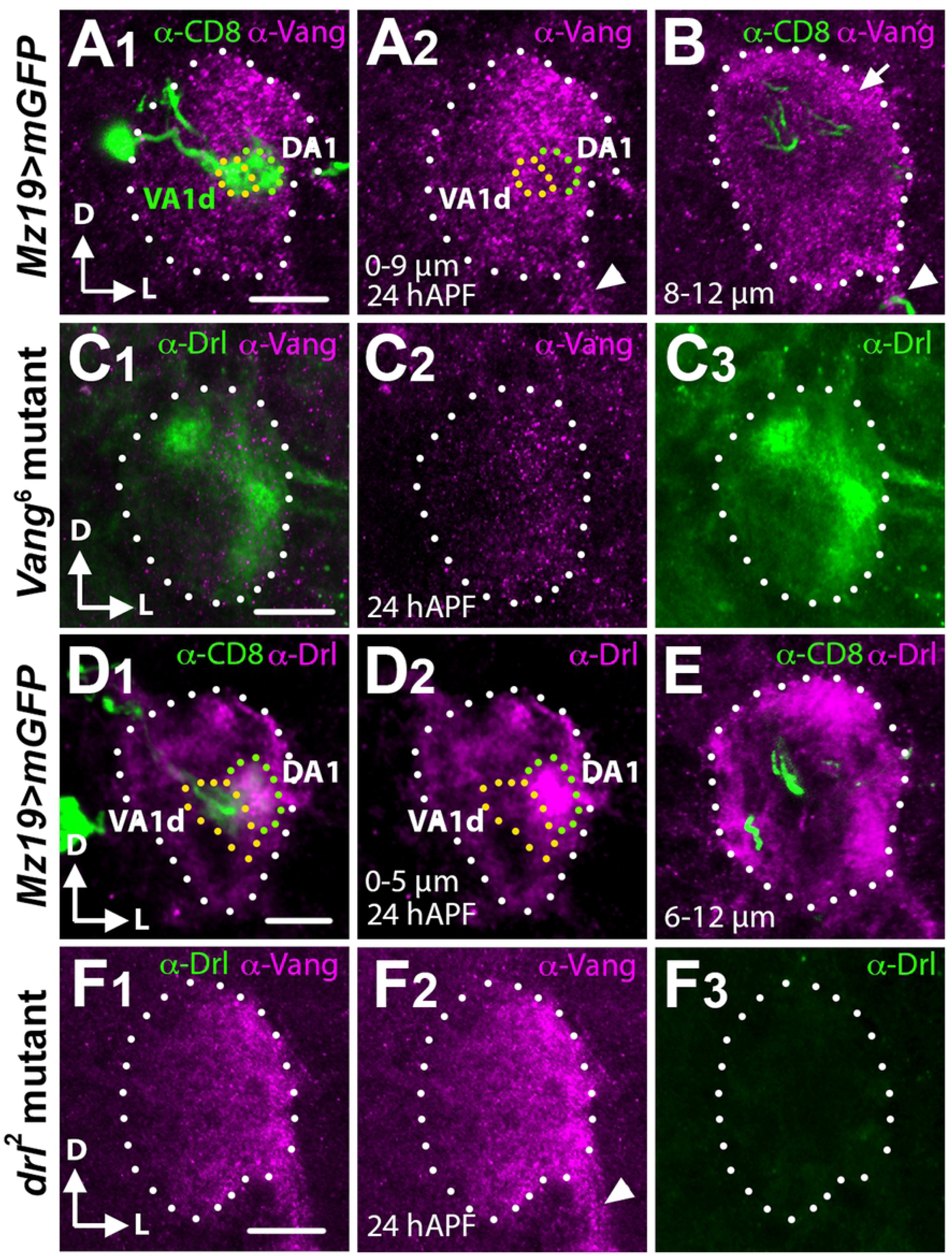
The Vang and Drl expression domains overlap with the developing DA1 and VA1d dendrites. Frontal views of a 24 hAPF ALs stained with Vang and Drl antibodies. (**A, B**) An AL from animals expressing *UAS-mCD8::GFP* under the control of *Mz19-Gal4* stained with antibodies against Vang (Magenta) and CD8 (green) to highlight the DA1 and VA1d PN dendritic arbors. Between 0-9 µm from the anterior AL surface Vang is found in puncta, which are highly concentrated in the AL dorsolateral region where the DA1 and VA1d dendrites are localized (A_1_, A_2_). In deeper sections (8-12 µm) Vang is observed in the nerve fiber layer (arrow) and the antennal nerve (arrowhead) (B). (**C**) An AL from *Vang^6^* mutants stained with antibodies against Vang (Magenta) and Drl (green) (C_1_). The AL stained poorly for Vang (C_2_) but strongly for Drl (C_3_) attesting to the specificity of the Vang antibody. (**D**, **E**) An AL from animals expressing *UAS-mCD8::GFP* under the control of *Mz19-Gal4* stained with antibodies against Drl (Magenta) and CD8 (green). From 0-5 µm the Drl protein is concentrated in the lateral AL where it co-localizes with the DA1 dendrites (D_1_, D_2_). The VA1d dendrites are located medially and express a low level of the Drl protein. Deeper down (5-12 µm) the Drl protein is found in the dorsal and ventrolateral neuropil structures (E). (**F**) An AL from *drl^2^* mutants stained with antibodies against Vang (Magenta) and Drl (green) (F_1_). The AL stained strongly for Vang (F_2_) but not at all for Drl (F_3_) attesting to the specificity of the Drl antibody. Scale bars: 10 µm.

### *Vang* is required in the ORNs for DA1-VA1d dendritic rotation

Since the Vang antibody strongly stained the AL nerve fiber layer, the antennal nerve and the antennal commissure, we hypothesized that Vang is expressed by ORNs and carried by axons to the developing AL. To identify the cell type in which *Vang* functions, we first used transgenic techniques to modulate *Vang* activity in specific cell types and examined the effect on the DA1/VA1d dendritic rotation. When we expressed the *UAS-Vang* transgene with the *Elav-Gal4* pan-neuronal driver (25) in the *Vang^6^* mutant, the DA1/VA1d dendritic angles became smaller (32.88° ± 2.19°, N=24) compared with that of the mutant control (51.40° ± 3.39°, N=20, *t*-test p<0.0001), indicating that *Vang* functions in neurons to promote dendritic rotation (Fig 3A-C, M). When we expressed *UAS-Vang* using the ORN-specific drivers, *peb-Gal4* and *SG18.1-Gal4*, the DA1/VA1d dendritic angles were also reduced (29.18° ± 1.6°, N=37 and 32.12° ± 2.12°, N=58 respectively) compared with that of the *Vang^6^* mutant (60.82° ± 2.52°, N=17, *peb* rescue vs. *Vang^6^*, p<0.0001; *SG18.1* rescue vs. *Vang^6^*, p<0.0001; *peb* rescue vs *SG18.1* rescue, p=0.8283; one-way ANOVA with post hoc Tukey test), indicating that *Vang* acts in the ORNs (Fig 3D-I, M). In contrast, expression of *UAS-Vang* using a PN-specific driver, *GH146-Gal4*, did not significantly alter the DA1/VA1d rotational angles of the *Vang^6^* mutant (47.87° ± 2.21°, N=39, *t*-test p=0.3894; Fig 3M). In further support of *Vang* acting in the ORNs, when we drove the *UAS-Vang^RNAi^* transgene (26) in the *Vang^6^/+* heterozygote with *peb-Gal4*, the rotation of the DA1/VA1d dendritic angle increased slightly compared with that of the *Vang^6^/+* heterozygote, indicating that *Vang* is required in the ORNs for DA1/VA1d dendritic rotation (36.40° ± 2.93°, N=24 compared with 30.09° ± 1.01°, N=22, *t*-test p=0.0518) (Fig 3J-L, N).

**Fig 3.**
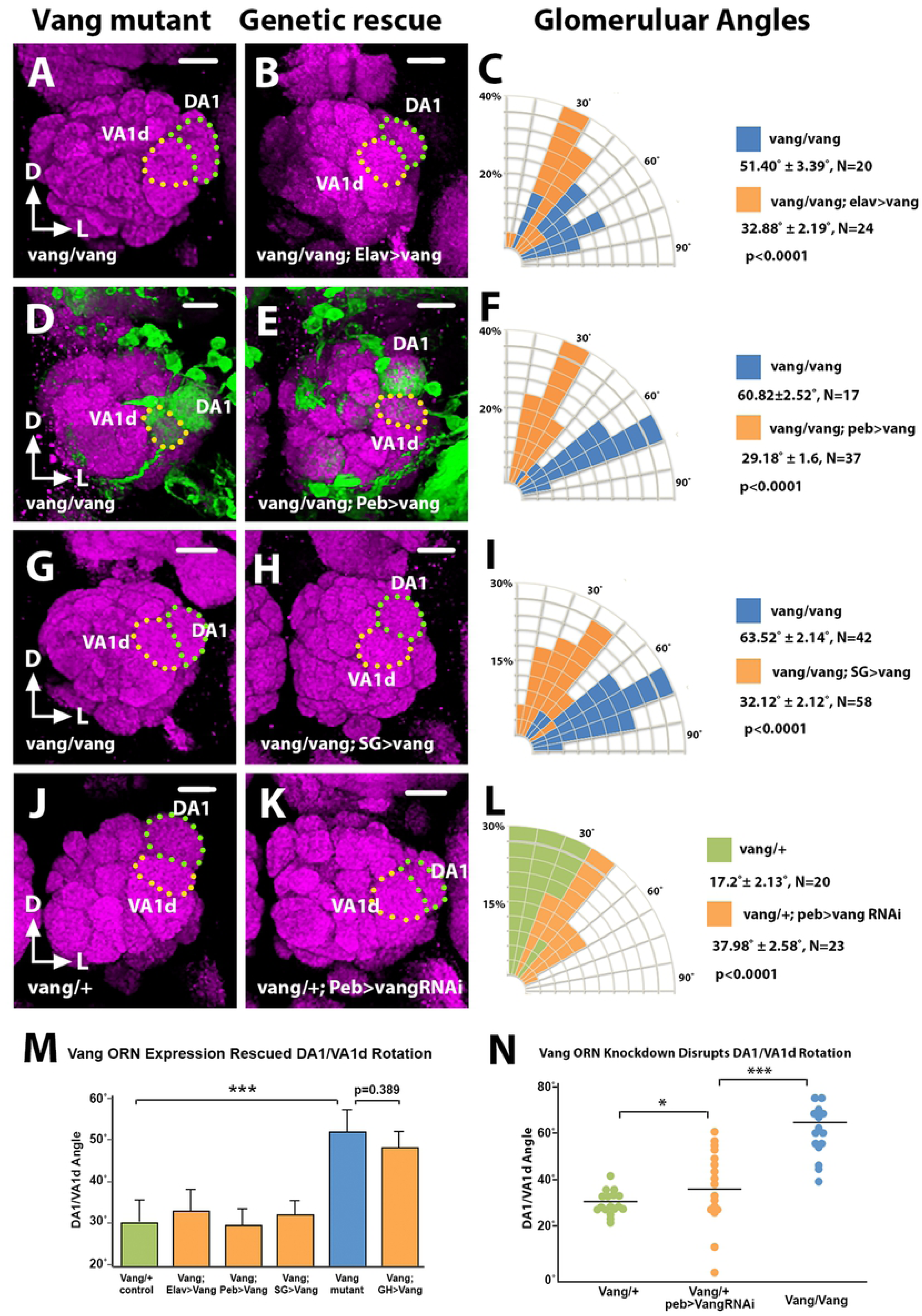
*Vang* functions in the ORNs to regulate glomerular migration. Adult *Vang^6^* ALs expressing *Mz19-mCD8::GFP* and different *Vang* transgenes were stained with antibodies against Brp (Magenta) and CD8 (green) to visualize the glomerular pattern. (**A, B**) Expression of *UAS-Vang* under the control of the pan-neuronal driver *Elav-Gal4* in the *Vang^6^* mutant (A) caused DA1 to migrate dorsally relative to the VA1d glomerulus (B). (**C**) Quantification of the DA1/VA1d glomerular angles in A and B. (**D, E, G, H**) Expression of *UAS-Vang* under the control of the ORN-specific drivers, *peb-Gal4* (D, E) and *SG18.1-Gal4* (G, H), also rescued DA1 dorsal migration. (**F, I**) Quantification of the DA1VA1d glomerular angles in D, E, G, and H. (**J, K**) Expression of *UAS-Vang^RNAi^* under the control of the *peb-Gal4* in the *Vang^6^/+* mutant (J) slightly disrupted DA1 dorsal migration (K). (**L**) Quantification of the DA1VA1d glomerular angles in J and K. (**M**) Graph summarizing the glomerular angles in the rescue experiments. ANOVAs were used to simultaneously compare the *Vang^6^* control and the rescued conditions. (**N**) Graph summarizing the glomerular angles in the *Vang^RNAi^* knockdown experiment. With the exception of M, Student’s *t* tests were used to compare the data of the mutants with those of their respective controls. Scale bars: 10 µm.

To confirm the above findings, we used mosaic techniques to induce ORNs or PNs lacking the *Vang* gene and examined the effects on the DA1/VA1d dendritic rotation. Induction of either *Vang^f04290^* or *Vang^6^* mutant ORN axons using the *ey-FLP/FRT* technique, which induces large clones in the antenna (27), resulted in the DA1/VA1d dendritic pair exhibiting larger angles (54.73° ± 3.26°, N=30 and 51.03° ± 2.62°, N=28 respectively) compared with animals innervated by wild-type ORN axons (20.24° ± 2.51°, N=20, wild type vs. *Vang^f04290^*, p<0.0001; wild type vs. *Vang^6^*, p<0.0001; *Vang^6^* vs *Vang^f04290^*, p=0.7297; one-way ANOVA with post hoc Tukey test) (Fig 4A-C, G, H), confirming that *Vang* is required in the ORNs for DA1/VA1d dendritic rotation. Next, we induced *Vang^6^* mutant PN clones using the MARCM system (28) with *GH146-Gal4* as the PN marker. We observed that *Vang^6^* mutant PN neuroblast and single-cell clones extended their dendrites into AL and innervated the glomeruli normally (Fig 4D-F). Importantly, ALs innervated by large *vang^6^* PN clones exhibited normal dendritic pattern, as judged by the angles of the DA1/VA1d dendrites (27.50° ± 4.89°, N=12) compared with those of the control (23.29° ± 5.27°, N=7, *t*-test p=0.56) (Fig 4E, G, H). Thus, our transgenic rescue and mosaic experiments showed that *Vang* functions in the ORNs to non-autonomously promote the rotation of the DA1/VA1d dendritic pair.

**Fig 4.**
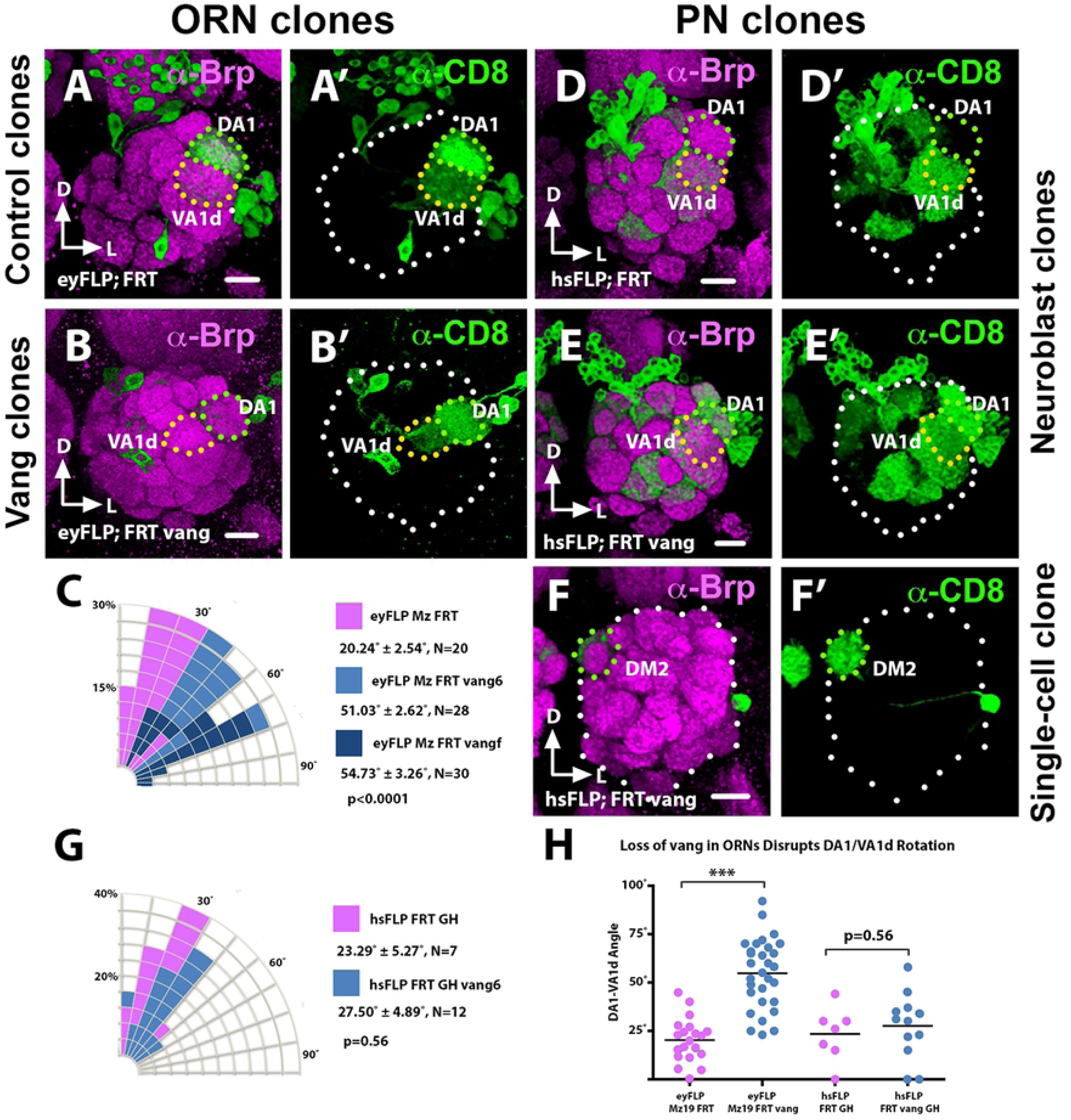
*Vang* functions selectively in ORNs but not the PNs to regulate glomerular migration. *Vang^6^* and control mosaic ALs were stained with antibodies against Brp (Magenta) to visualize the glomerular pattern and CD8 (green) to visualize PN dendritic arborization. (**A**) In ALs innervated by wild-type ORN axon clones induced by the *ey-FLP/FRT* technique the DA1 glomerulus migrated normally relative to the VA1d glomerulus. (**B**) In ALs innervated by *Vang^6^* mutant axon clones induced by the *ey-FLP/FRT* technique the DA1 glomerulus failed to migrate dorsally relative to the VA1d glomerulus. (**C**) Quantification of the glomerular angles in A and B. ANOVAs were used to simultaneously compare the control and the *Vang^6^* and *Vang^f04290^* conditions. (**D**) In ALs innervated by large clones of wild-type PN dendrites, induced by the MARCM technique the DA1 glomerulus migrated normally relative to the VA1d glomerulus. (**E, F**) In ALs innervated by neuroblast (E) and single-cell (F) clones of *Vang^6^* PN dendrites induced by the MARCM technique, the mutant dendrites innervated the AL normally and the DA1 glomerulus migrated normally relative to the VA1d glomerulus. (**G**) Quantification of the glomerular angles in D, E and F. Student’s *t* test was used to compare the data of the mutant clones with that of the controls. (**H**) Graph summarizing the glomerular angles in the mosaic control and *Vang^6^* ALs. Scale bars: 10 µm.

### *Vang* is not required for ORN axon growth or correct ORN-PN pairing

A possible explanation for the *Vang* mutant phenotype is that *Vang* is required for ORN axon growth to the AL, the failure of which indirectly disrupted glomerular patterning. Mutations in *Vang* have been shown to result in abnormal projection of mushroom body axons (29). To determine if ORN axons entered the AL in the *Vang^6^* mutant, we labeled eight different ORN axon terminals in the AL using *Or-Gal4* drivers. We found that *Vang^6^* mutant axons entered the AL normally, although their terminals were shifted in the AL neuropil (Fig 1 and data not shown). To investigate if the *Vang* mutation disrupted the proper matching of the ORN axons and PN dendrites, we simultaneously labeled pre- and postsynaptic partners of glomeruli for which specific markers were available. We achieved this by labeling the DA1, VA1d and DM1 dendrites with *Mz19-Gal4* driving *UAS-mCD8::GFP* and simultaneously the ORN axons targeting the VA1d and VA1lm glomeruli with the *Or88a-CD2* and *Or47b-CD2* transgenes respectively. We observed that *Or88a* axons were strictly paired with VA1d PN dendrites in the *Vang^6^* mutant as in the wild type (Fig 5A, B). Likewise, the *Or47b* axons strictly innervated the VA1lm glomerulus, and never strayed into the VA1d territory in the *Vang^6^* mutant (Fig 5C). Thus, *Vang* is not required for ORN axon projection into the AL or their correct pairing with their postsynaptic partners. We propose that *Vang* functions in the context of paired axons and dendrites allowing the neurites to coordinately respond to the *Wnt5* signal. This idea is consistent with our observation that PN dendritic rotation occurs between 16 and 30 hAPF (11), the period of major ORN axon invasion into the AL (8, 12).

**Fig 5.**
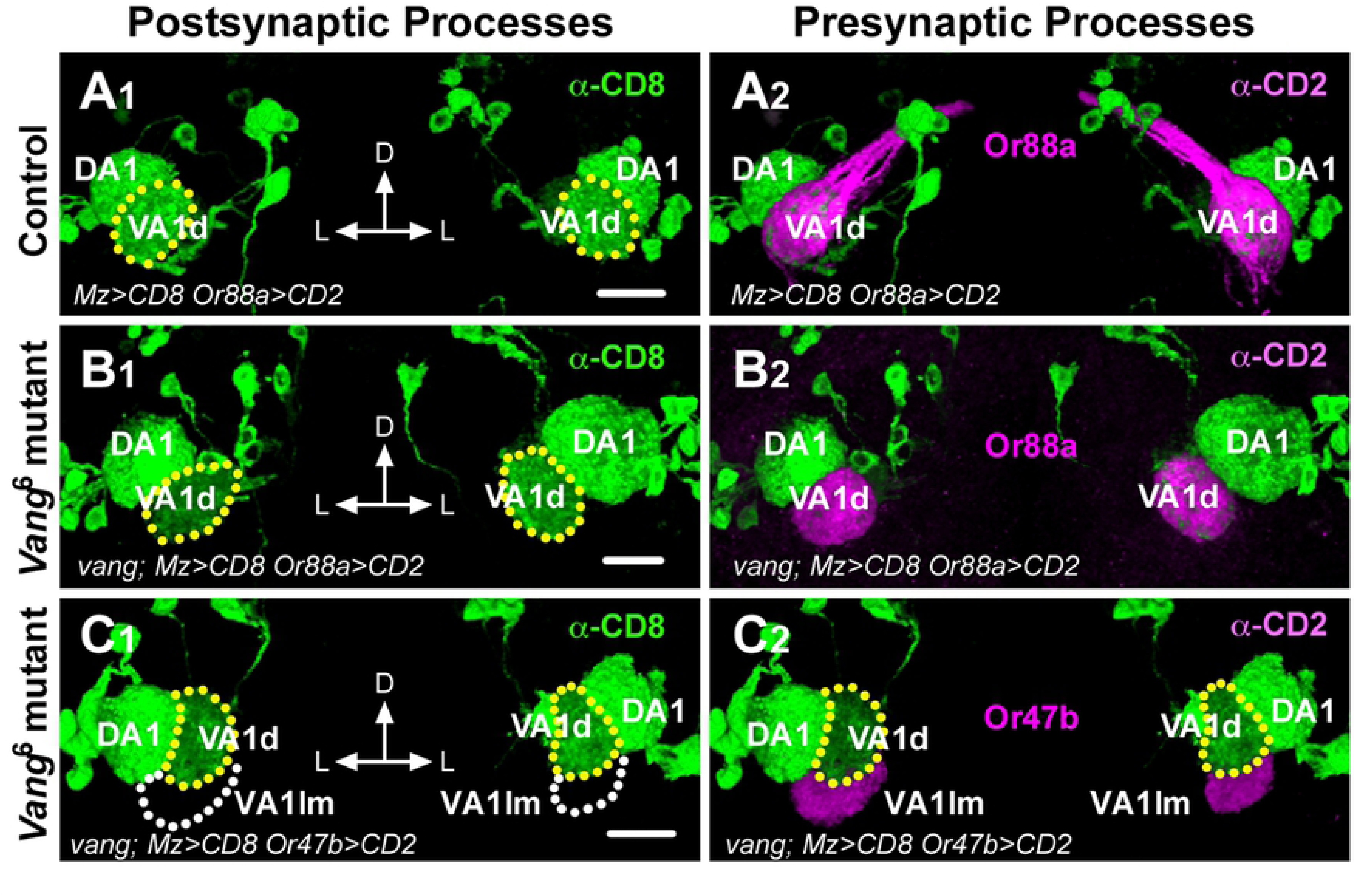
*Vang* does not function in ORN axon projection or pairing with cognate PN dendritic partners. (**A, B**) Adult brains from animals expressing *Or88a-CD2* and *UAS-mCD8::GFP* under the control of *Mz19-Gal4* were stained with anti-CD2 (magenta) and anti-CD8 (green) to visualize the pre- and postsynaptic processes of the VA1d and VA1lm glomeruli. In the wild-type control, *Or88a* axons are faithfully paired with the VA1d dendrites (A). Collaterals form a fascicle, which innervates the contralateral AL. In the *Vang^6^* mutant, *Or88a* axons are also faithfully paired with the VA1d dendrites (B). *Vang* mutant axons fail to sprout collaterals as previously reported (Shimizu). (**C**) Frontal views of adult ALs from *Vang^6^* mutants expressing *Or47b-CD2* and *UAS-mCD8::GFP* under the control of *Mz19-Gal4* stained with anti-CD2 (magenta) and anti-CD8 (green). In the mutant, *Or47b* axons targeted the adjacent VA1lm glomerulus without straying into the VA1d glomerular territory. Scale bar: 10 µm.

### *Vang* acts downstream of *Wnt5* to repel the VA1d glomerulus

The close resemblance of the *Vang* and *Wnt5* mutant phenotypes raised the questions of whether and how *Vang* might function in the *Wnt5* signaling pathway to regulate the rotation of the DA1 and VA1d glomeruli. To address these questions, we asked if loss of *Vang* would block *Wnt5* signaling. We previously showed that overexpression of *Wnt5* in the DA1 and VA1d dendrites with the *Mz19-Gal4* driver split the VA1d dendrites into two smaller arbors probably due to repulsion between the dendrites (Fig 6A, B) (11). Interestingly, *Wnt5* overexpression had no effect on the DA1 dendrites, indicating that the DA1 and VA1d dendrites respond differentially to the *Wnt5* signal. The VA1d defect provided an opportunity to assess if *Vang* is needed for the *Wnt5* gain-of-function phenotype. Whereas only 9.37% (3/32) of the VA1d dendrites in the *Mz19>Wnt5* animals were intact, this fraction rose to 45.65% (21/46) in the *Mz19>Wnt5; Vang^6^/Vang^6^* animals (Fig 6C, F). Moreover, the distances between the split VA1d arbors in the *Mz19>Wnt5; Vang^6^/Vang^6^* animals were smaller than those in the *Mz19>Wnt5* animals (11.83 µm ± 0.61 µm, N=25, vs 21.06 µm ± 0.96 µm, N=29, *t*-test p<0.0001) (Figs. 6B, C, G). Despite severe distortion, the VA1d dendrites were faithfully paired with their *Or88a* axon partners, reinforcing the idea that *Wnt5* signaling does not play a role in ORN-PN matching (Fig 6D-E). We conclude that *Wnt5* signals through *Vang* to repel the VA1d glomerulus.

**Fig 6.**
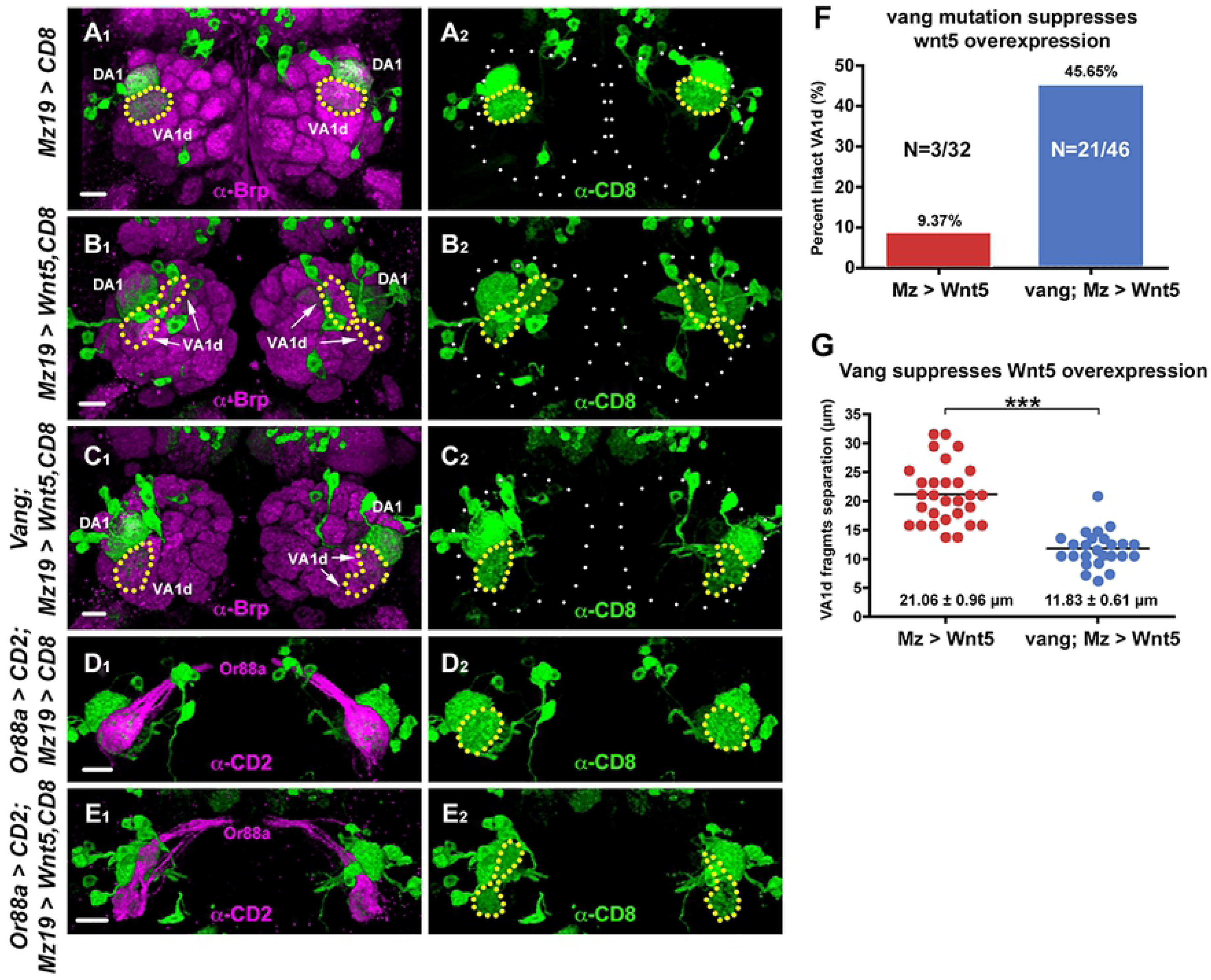
*Vang* functions downstream of *Wnt5* to cell non-autonomously repel the VA1d dendrites. (**A-C**) Adult ALs from animals expressing *UAS-mCD8::GFP* under the control of *Mz19-Gal4* were stained with antibodies against Brp (magenta) to visualize the glomerular pattern and CD8 to visualize the DA1/VA1d dendrites (green). (A) In the wild-type control, the DA1 dendrites are located dorsal to the VA1d dendrites, which form a single compact arbor. (B) In animals expressing *UAS-Wnt5* under the control of *Mz19-Gal4*, the VA1d dendrites split into two separated arbors, probably due to repulsion between the two dendritic branches. (C) Removing *Vang* functions in animals expressing *UAS-Wnt5* under the control of *Mz19-Gal4* return the VA1d dendrites to its compact morphology in the right AL. In the left AL, the two separated branches are closer together. (**D, E**) Adult ALs from animals expressing *Or88a-CD2* and *UAS-mCD8::GFP* under the control of *Mz19-Gal4* were stained with anti-CD2 (magenta) and anti-CD8 (green) to visualize the pre- and postsynaptic processes of the VA1d glomerulus. In the wild-type control, *Or88a* axons are faithfully paired with the VA1d dendrites (D). In animals expressing *UAS-Wnt5* under the control of *Mz19-Gal4* the *Or88a* axons were still correctly paired with the VA1d dendrites despite their splitting into two separated arbors (E). (**F**) Graph summarizing the percentage of intact VA1d glomeruli in wild-type and *Vang^6^* animals overexpressing *Wnt5* in the VA1d glomeruli. (**G**) Graph summarizing the distance between the split VA1d arbors in wild-type and *Vang^6^* animals overexpressing *Wnt5* in the VA1d glomeruli. Student’s *t* test was used to compare the data of the wild-type and *Vang^6^* animals overexpressing *Wnt5.* Scale bars: 10 µm.

To further probe the relationship between *Wnt5* and *Vang*, we examined the DA1/VA1d rotation in animals lacking both genes. We observed that the rotation in the *Wnt5^400^; Vang^6^* double mutant (92.20° ± 4.1°, N=41) is more severely disrupted than that in either single mutant (76.03° ± 3.62° in *Wnt5^400^*, n=29, *t*-test p=0.0031 and 54.12° ± 2.8° in *Vang^6^*, n=34, *t*-test p<0.0001) (Fig 7). The enhanced phenotype of the *Wnt5^400^; Vang^6^* mutant suggested that *Wnt5* and *Vang* could function independently to promote DA1/VA1d rotation (Fig 7F). We currently do not know how *Vang* acts independently of *Wnt5*. However, it interesting that *Wnt5* directs the rotation of the DA1/VA1d glomeruli through both *Vang*-dependent and *Vang*-independent pathways. Since *Wnt5* acts through *Vang* in the VA1d glomerulus, we posited that *Wnt5* acts through a *Vang*-independent mechanism in the DA1 glomerulus.

**Fig 7.**
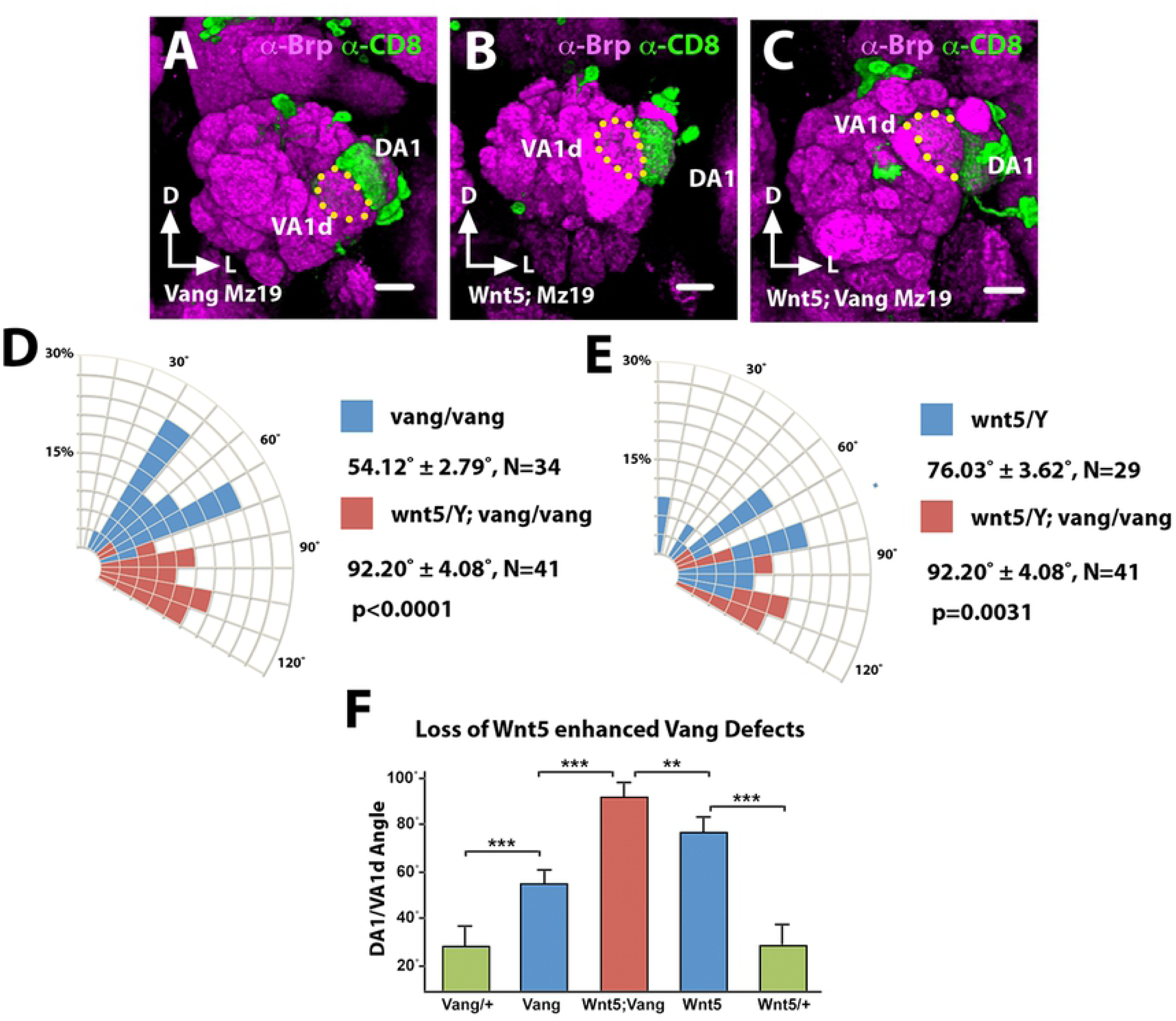
*Wnt5* and *Vang* could function independently to regulate DA1/VA1d glomerular rotation. (**A-C**) Frontal views of adult ALs from animals expressing *UAS-mCD8::GFP* under the control of *Mz19-Gal4* stained with antibodies against Brp (magenta) and CD8 (green) to visualize the glomerular pattern and the DA1/VA1d dendrites respectively, in the *Vang^6^* (A), *Wnt5^400^* (B) and *Wnt5^400^; Vang^6^* (C) mutants. (**D**) Quantification of the DA1/VA1d angles in panels A and C showing that the rotational defect is stronger in the *Wnt5^400^; Vang^6^* mutant than in the *Vang^6^* mutant. (**E**) Quantification of the DA1/VA1d angles in B and C showing that the rotational defect is stronger in the *Wnt5^400^; Vang^6^* mutant than in the *Wnt5^400^* mutant. (**F**). Graph summarizing the DA1/VA1d angles in the *Vang^6^* and *Wnt5^400^* single mutants and *Wnt5^400^; Vang^6^* double mutant. Student’s *t* tests were used to compare the angles of the various mutants. Scale bars: 10 µm.

### *drl* promotes the rotation of DA1-VA1d glomeruli

The Drl atypical receptor tyrosine kinase has been shown to bind Wnt5 and mediates its signaling in the migration of a number of cell types (14–17). We previously demonstrated that Drl is differentially expressed by PN dendrites wherein it antagonizes Wnt5’s repulsion of the dendrites (11). To delineate the AL region where *drl* functions, we examined the positioning of several glomeruli in the null *drl^2^* mutant by expressing *UAS-mCD8::GFP* under the control of various *Or-Gal4* drivers (Fig 8A-I). We observed that, as in the *Vang* mutant, the lateral glomeruli showed the strongest displacements in positions compared with the control indicating that *drl* primarily regulates neurite targeting in the lateral AL (Fig 8A-C). To better characterize the neuropil defect in this region we employed the *Crispr/Cas9* technique (30) to create the null *drl^JS^* allele on the *Mz19-Gal4* chromosome (Materials and Methods), which allowed us to assess the DA1/VA1d dendritic arrangement in the *drl* mutant. To our surprise, we observed that the DA1/VA1d dendritic pair in the *drl^JS^*/*drl^2^* null mutant showed strong deficits in DA1/VA1d rotation, resembling the *Wnt5^400^* null phenotype (Fig 8J-L). Indeed, measurement of the DA1/VA1d angle of the *drl^JS^*/*drl^2^* mutant (76.55° ± 3.75°, N=31) showed that it was even slightly larger than that of the *Wnt5^400^* mutant (69.22° ± 5.03°, N=32, *t*-test p=0.2493) (Fig 10G). The similarity of the *drl* and *Wnt5* mutant phenotypes indicates that *drl* cooperates with *Wnt5* in promoting the rotation of the DA1/VA1d glomeruli.

**Fig 8.**
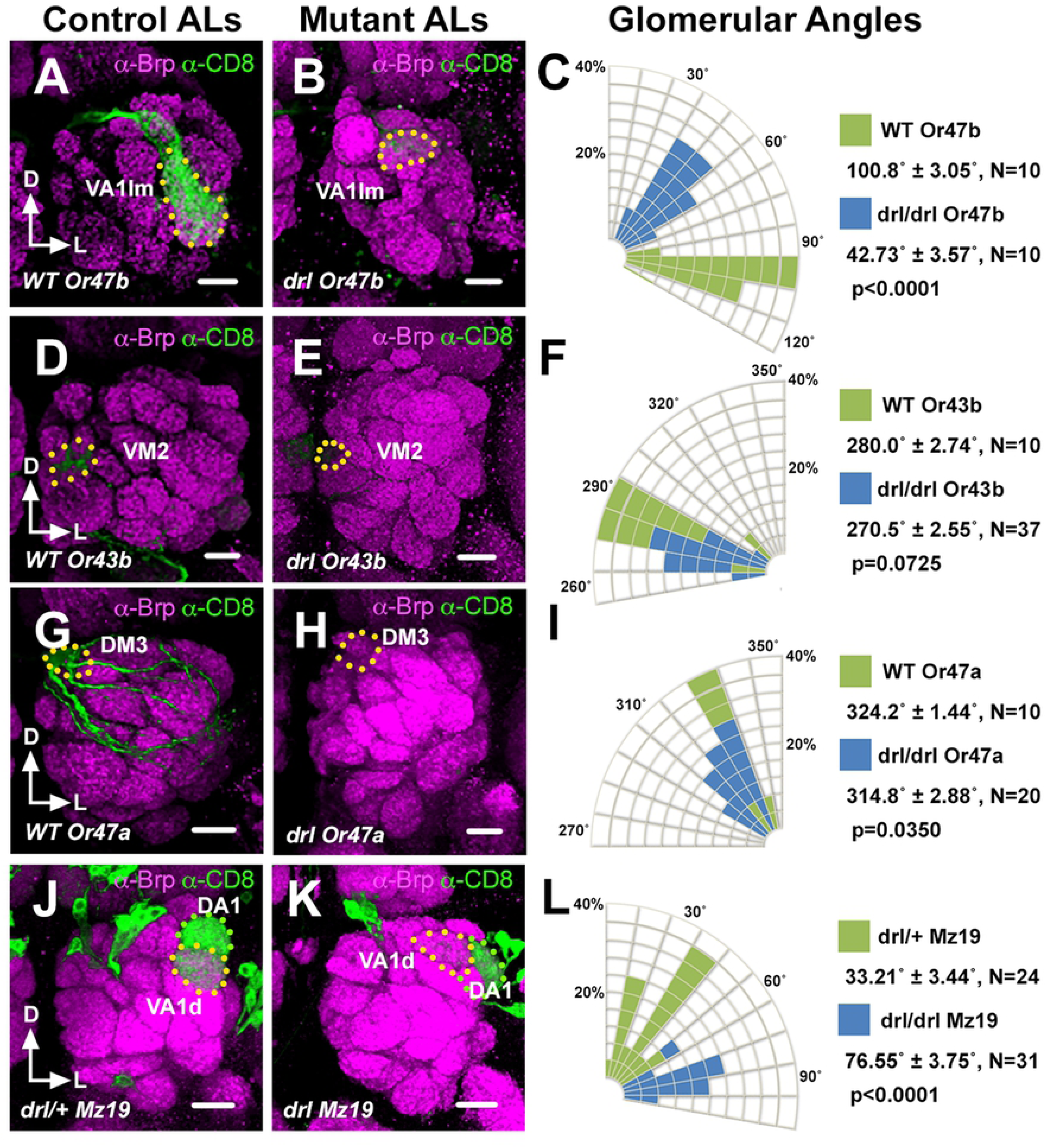
The *drl^2^* mutant AL defects resemble that of the *wnt5^400^* mutant. (**A-I**) Left adult ALs from wild-type (A, B, G) and *drl^2^* (B, E, H) animals expressing *UAS-mCD8::GFP* under the control of *Or47b-Gal4* (A, B), *Or43b-Gal4* (D, E) and *Or47a-Gal4* (G, H) were stained with antibodies against Brp (Magenta) to highlight the AL neuropil and CD8 (green) to label lateral, medial and dorsal glomeruli respectively. (**C, F, I**) Quantification of the positions of the *Or47b*, *Or43b* and *Or47a* glomeruli respectively in the wild-type vs *drl^2^* animals. The glomeruli appeared to be displaced in the counterclockwise direction in the *drl^2^* mutant with VA1lm showing the greatest displacement. (**J, K**) ALs from adult *drl^JS^/+* control (J) and *drl^JS^/drl^2^* animals (K) expressing *UAS-mCD8::GFP* under the control of *Mz19-Gal4* were stained with antibodies against Brp (Magenta) and CD8 (green) to highlight the AL neuropil and the DA1/VA1d PN dendritic arbors respectively. (**L**) Quantification of the DA1/VA1d angles in the *drl^JS^/drl^2^* mutant. The DA1 dendrites are located lateral to the VA1d dendrites in the *drl^JS^/drl^2^* mutant reflecting a severe impairment in DA1/VA1d dendritic rotation. This phenotype resembles that of the *Wnt5^400^* mutant. Student’s *t* tests were used to compare the data of the *drl* mutants with those of the controls. Scale bars: 10 µm.

### *drl* acts in the DA1 dendrites to promote glomerular attraction to *Wnt5*

How does *drl* mediate *Wnt5* function in glomerular rotation? We hypothesized that *drl* functions in the DA1 dendrites, to regulate migration of the DA1 glomerulus towards the *Wnt5* source. In support of this idea, antibody staining of the Drl protein showed that it is highly expressed by the DA1 dendrites but not the VA1d dendrites (Fig 2D, E) (11). The domain of high Drl expression occupies the anterior dorsolateral domain of the 24 hAPF ALs (0-8 µm from anterior), a region in which Wnt5 is also highly expressed (11). The hypothesis predicts that ablation of *drl* in DA1 alone would disrupt the rotation of the DA1/VA1d glomeruli. To test this hypothesis, we used MARCM to induce *drl^JS^* homozygosity in either DA1 or VA1d dendrites and assessed the effects on DA1/VA1d glomerular rotation. We observed that *drl^JS^* mutant VA1d dendrites were associated with glomerular pairs with small angles (17.13° ± 3.68°, N=16), that is, with the DA1 glomerulus closely associated with the dorsolateral AL (Fig 9C-F). In contrast, *drl^JS^* mutant DA1 dendrites are associated with glomeruli with wide variations in angles (58.43° ± 8.33°, N=14, *t*-test p=0.0003), that is, with the DA1 glomerulus not closely associated with the dorsolateral AL (Fig 9A, B, E, F). Thus, *drl* appears to act in the DA1 dendrites to confer directionality of the DA1 glomerulus towards the *Wnt5* source.

**Fig 9.**
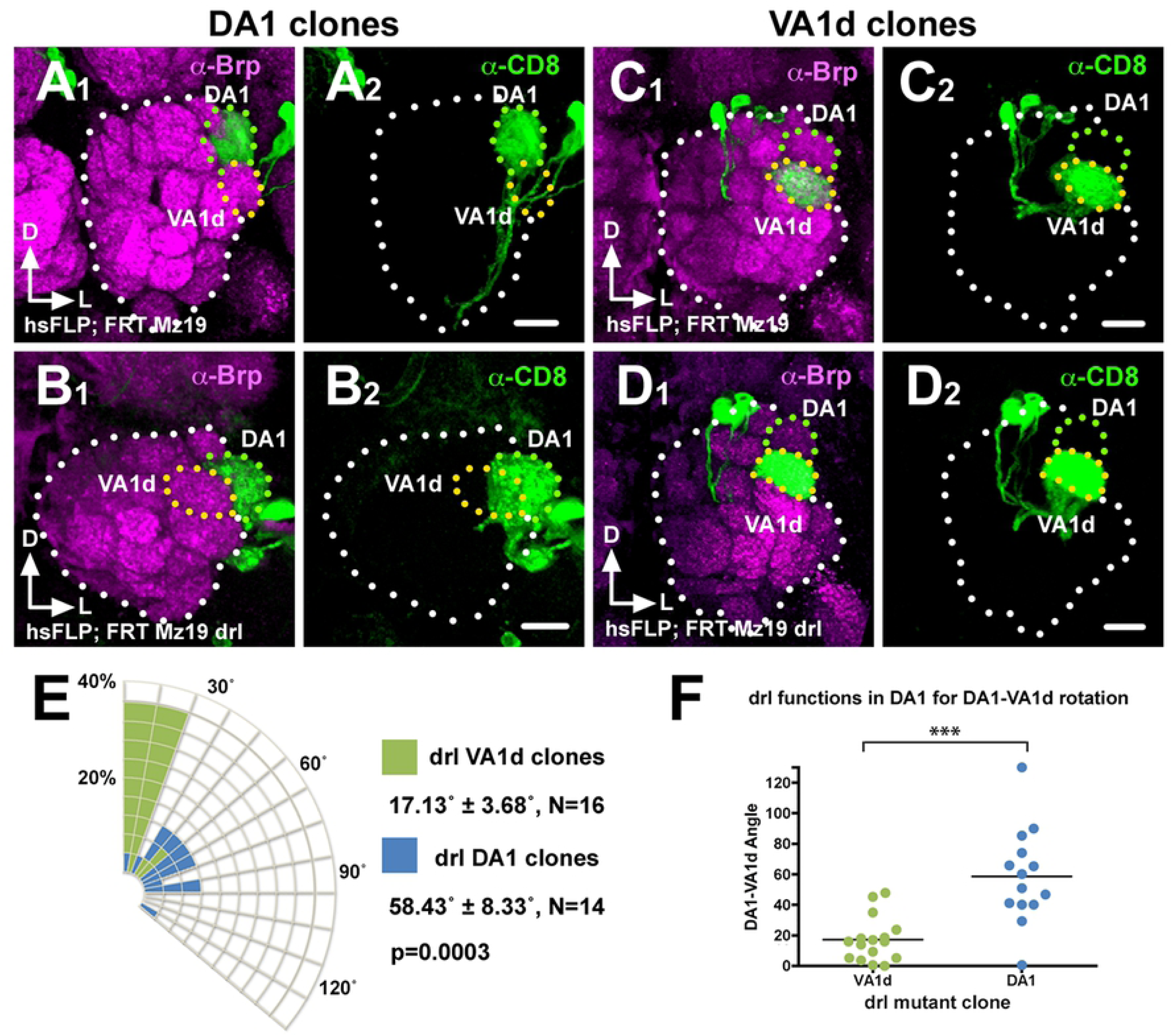
*drl* functions in the DA1 dendrites but not VA1d dendrites to promote DA1/VA1d glomerular rotation. Mosaic control and *drl^JS^* ALs generated by the MARCM technique were stained with antibodies against Brp (Magenta) to visualize the glomerular pattern and CD8 (green) to visualize the clonal PN dendrites. (**A**) Control DA1 dendrites are often observed in DA1/VA1d glomerular pair with small rotational angles. (**B**) In contrast, *drl^JS^* mutant DA1 dendrites are frequently observed in DA1/VA1d glomeruli with large rotational angles indicating impaired glomerular rotation. (**C**) Control VA1d dendrites are often observed in DA1/VA1d glomerular pair with small rotational angles. (**D**) Likewise, *drl^JS^* mutant VA1d dendrites are frequently observed in DA1/VA1d glomerular pair with small rotational angles indicating normal glomerular rotation. (**E**) Quantification of the glomerular angles of the DA1 and VA1d clones. (**F**) Graph summarizing the *drl^JS^* DA1 vs VA1d clonal data. Student’s *t* test was used to compare the data of the *drl* DA1 vs VA1d mutant clones. Scale bars: 10 µm.

**Fig 10.**
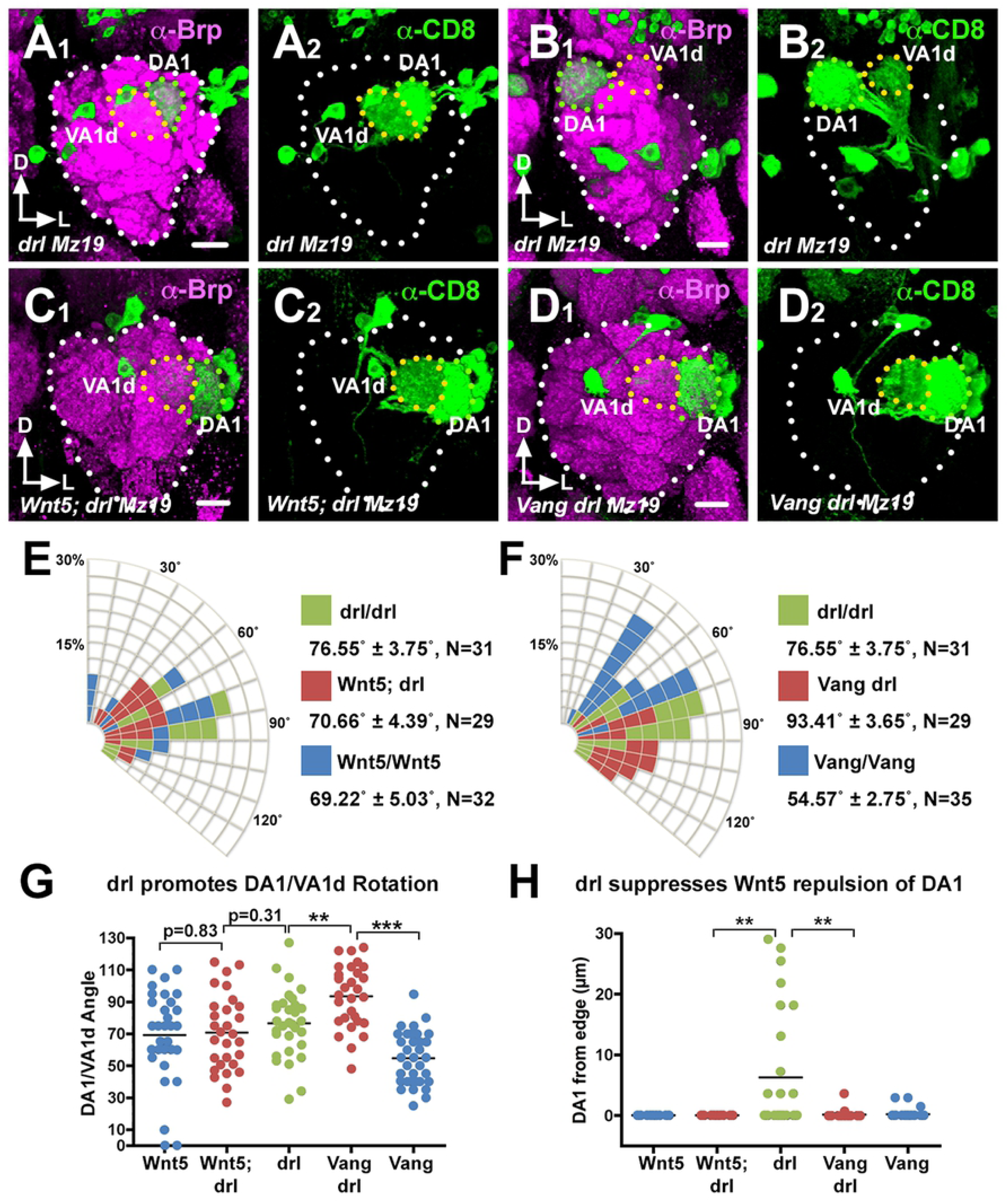
*Wnt5* repels the DA1 glomerulus through *Vang*, a function that *drl* antagonizes. Left adult ALs of animals expressing *UAS-mCD8::GFP* under the control of *Mz19-Gal4* were stained with nc82 (magenta) and anti-CD8 (green) to visualize the neuropil and the DA1/VA1d dendrites respectively. (**A, B**) In the *drl* mutant, the DA1 glomerulus was often displaced from the AL lateral border, resulting in the medial shift of the DA1/VA1d glomerular pair (A) or a reversal in the positions of the two glomeruli (B). (**C**) The loss of the *wnt5* gene suppressed the DA1 medial displacement, restoring the DA1 glomerulus to the AL lateral border, indicating that *drl* antagonizes *Wnt5* repulsion of the DA1 glomerulus. (**D**) The DA1 medial displacement is also suppressed by the loss of the *Vang* gene, suggesting that *Vang* functions in *Wnt5* repulsion of the DA1 glomerulus. In addition, the DA1/VA1d glomerular angle of the *drl* mutant is large. (**E**) Quantification of the DA1/VA1d glomerular angles in the *drl* and *Wnt5* single mutants and *Wnt5; drl* double mutant. (**F**) Quantification of the DA1/VA1d glomerular angles in the *drl* and *Vang* single mutants and *Vang drl* double mutant. (**G**) Graph summarizing the DA1/VA1d glomerular angles in the *drl*, *Wnt5*, and *Vang* mutants. (**H**) Graph summarizing the distance of the DA1 glomerulus from the AL lateral border in the *drl*, *Wnt5*, and *Vang* mutants. Student’s *t* tests were used to compare the data of the double mutants with the single mutants. Scale bars: 10 µm.

### *drl* likely converts *Wnt5* repulsion of the DA1 glomerulus into attraction

How does *drl*, which antagonizes *Wnt5* signaling (11), promote the migration of the DA1 glomerulus towards *Wnt5*? We propose two models by which *drl* could accomplish this task. In the first model, *drl* acts as a positive effector of *Wnt5* attractive signaling. In the second model, *drl* neutralizes *Wnt5* repulsive signaling and/or converts the repulsive signaling into an attractive one. Careful examination of the DA1/VA1d targeting defects in *drl^JS^*/*drl^2^* null mutant revealed differences with that of the *Wnt5^400^* mutant, inconsistent with the idea that *drl* acts as a positive effector of *Wnt5* signaling. First, the DA1 glomerulus is often displaced medially from the AL lateral border (6.26 µm ± 1.85 µm from border, N=28) (Fig 10H), a defect not seen in the *Wnt5^400^* mutant. This resulted in the frequent reversal in the positions of the DA1 and VA1d glomeruli (Fig 10B), or displacement of both glomeruli medially from the AL lateral border (Fig 10A). Second, the mean DA1/VA1d angle in the *drl^JS^*/*drl^2^* null mutant is slightly but not significantly larger than that of *Wnt5^400^* null mutant (Fig 10E, G). Instead the defects are more consistent with the second model, which predicts increased *Wnt5* repulsion of the DA1 glomerulus in the *drl* mutant, thus driving the glomerulus ventromedially. To test this hypothesis, we simultaneously removed both *drl* and *Wnt5* functions and examined the displacement of the DA1/VA1d glomeruli. We observed that the DA1 glomerulus is restored to AL lateral border in the *Wnt5^400^*; *drl^JS^*/*drl^2^* double mutant, as it is in the *Wnt5* homozygote (0.00 µm from edge, N=30 and N=25 respectively) (Fig 10C, H). We also observed that the DA1/VA1d angle in the *Wnt5; drl* double mutant (70.66° ± 4.39°, N=29) is more similar to that of the *Wnt5* mutant (69.22° ± 5.03°, N=32, *t*-test p=0.83) than that of the *drl* mutant (76.55° ± 3.75°, N=31, *t*-test p=0.3102) (Fig 10E, G). We conclude that *Wnt5* repels the DA1 glomerulus ventromedially and that *drl* antagonizes the *Wnt5* repulsive activity. Since we showed above that *drl* promotes the migration of DA1 towards *Wnt5*, we conclude that *drl* acts in the DA1 glomerulus to convert *Wnt5* repulsion of the DA1 glomerulus into attraction. Taken together, our results suggest that *Wnt5* orients the rotation of the VA1d/DA1 glomeruli by attracting the DA1 glomerulus through *Wnt5-drl* signaling and repelling the VA1d glomerulus through *Wnt5-Vang* signaling.

To investigate the mechanism by which *drl* converts *Wnt5* repulsion of the DA1 glomerulus into attraction, we examined the mechanism by which *Wnt5* repels the DA1 glomerulus. A likely scenario is that *Wnt5* repels the DA1 glomerulus through *Vang*. To test this idea, we simultaneously removed both *drl* and *Vang* functions and measured the displacement of the DA1 glomerulus from the AL lateral border as well as the DA1/VA1d rotational angle. We found that in the *Vang^6^ drl^JS^*/*Vang^6^ drl^2^* double mutant, the DA1 glomerulus is restored to the AL lateral border (0.156 µm ± 0.132 µm, N=28, *t*-test p=0.0017) (Fig 10D, H). We conclude that *drl* neutralizes the *Wnt5-Vang* repulsion of the DA1 glomerulus. Interestingly, measurement of the DA1/VA1d angles in the *Vang drl* double mutant showed that the rotation of the glomeruli is more severely impaired (93.41° ± 3.65° N=29) than that of either single mutants (76.55° ± 3.75° in *drl^2^*, N=31, *t*-test p=0.0021 and 54.57° ± 2.75° in *Vang^6^*, N=35, *t*-test p<0.0001) (Fig 10F, G). The enhanced phenotype of the *Vang drl* double mutant indicated that *drl* and *Vang* act in parallel pathways to promote DA1/VA1d rotation. The parallel functions are in accord with our model that *drl* acts in the DA1 glomerulus while *Vang* acts in the adjacent VA1d glomerulus to promote DA1/VA1d rotation. Simultaneous loss of *Vang* and *drl* would be expected to exacerbate the DA1/VA1d rotational defect.

## Discussion

Elucidating the mechanisms that shape dendritic arbors is key to understanding the principles of nervous system assembly. Genetic approaches have revealed both intrinsic and extrinsic cues that regulate the patterning of dendritic arbors (31, 32). In contrast, there are only a few reports on the roles of axons in shaping dendritic arborization (33, 34). In this paper we provide evidence that final patterning of the fly olfactory map is the result of an interplay between ORN axons, PN dendrites and the Wnt5 directional signal (Fig 11). We show that the Vang PCP protein (20, 21) is an axon-derived factor that mediates the Wnt5 repulsion of the VA1d dendrites. We also show that the Drl protein is specifically expressed by the DA1 dendrites where it antagonizes the Wnt5-Vang repulsion of the DA1 glomerulus and likely converts it into an attractive response. The differential responses of the DA1 and VA1d glomeruli to Wnt5 would produce the forces by which Wnt5 effects the rotation of the glomerular pair. We present the following lines of evidence in support of this model of olfactory neural circuit development.

**Figure 11.**
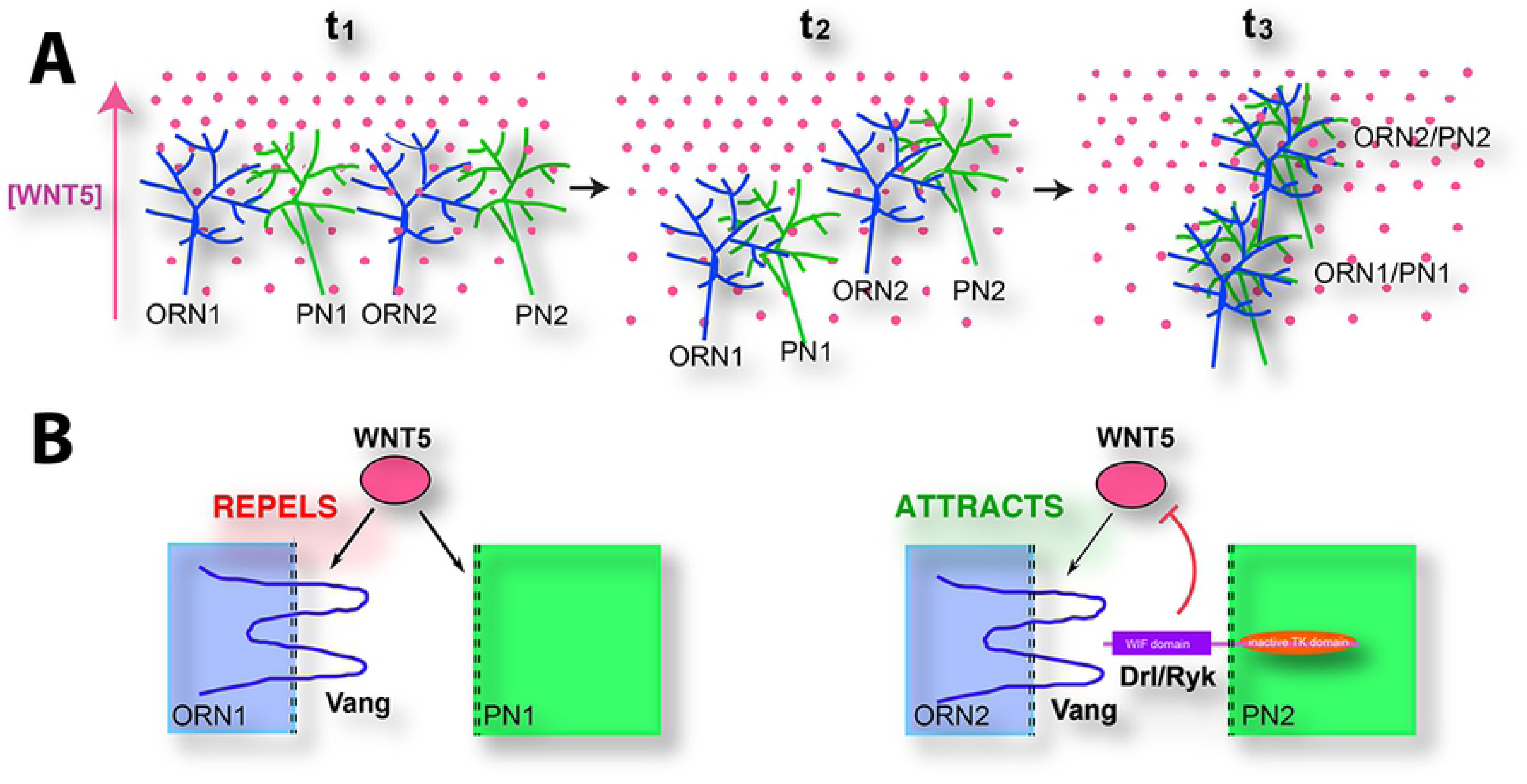
Model for the Wnt5-directed rotation of developing glomeruli. (**A**) The targeting of two ORN axons (ORN1 and ORN2) on their respective PN dendritic partners (PN1 and PN2) in the Wnt5 gradient at three time points is depicted. At time t_1_, the pre- and postsynaptic processes have not paired up and the individual neurites do not respond to the Wnt5 signal. At time t_2_, the neurites are beginning to pair up, which allows the nascent synapses to respond to the Wnt5 signal. (**B**) The ORN1:PN1 glomerulus is repelled by Wnt5 because the PN1 dendrite does not express Drl. On the other hand, the ORN2:PN2 glomerulus is attracted by Wnt5 because the PN2 dendrite expresses Drl, which converts Wnt5-Vang repulsion into attraction. Repulsion of the ORN1:PN1 glomerulus causes it to move down the Wnt5 gradient while the attraction of the ORN2:PN2 glomerulus causes it to move up the Wnt5 gradient. At time t_3_, the ORN axons and PN dendrites have fully condensed to form glomeruli. The opposing responses of the glomeruli to Wnt5 result in their rotation around each other.

Immunostaining showed that Vang is expressed at the same time and place as Wnt5 and Drl, and concentrated in the dorsolateral AL where major dendritic reorganization occurs (11). Mutations in *Vang* strongly disrupted the pattern of glomeruli in the AL, mimicking the *Wnt5* mutant phenotype. Notably, mutation of *Vang* suppressed the strong repulsion of the VA1d dendritic arbor caused by *Wnt5* overexpression, indicating that *Vang* acts downstream of *Wnt5* to repel the VA1d dendrites. Unexpectedly, using cell type-specific transgenic experiments and mosaic analyses we found that *Vang* functions specifically in the ORNs, indicating an obligatory codependence of ORN axon and PN dendritic targeting. Unlike the VA1d glomerulus, which expresses low levels of *drl* and is repelled by *Wnt5*, the adjacent DA1 glomerulus expresses high levels of *drl*. Mosaic analyses showed that *drl* acts specifically in the DA1 dendrites to confer directionality of the DA1 glomerulus towards *Wnt5*. Finally in the absence of *drl*, the DA1 glomerulus is displaced away from the *Wnt5* source, a defect that is suppressed by the removal of either *Wnt5* or *Vang*. Taken together, we propose that *drl* likely converts *Wnt5* repulsion of the DA1 glomerulus into attraction by inhibiting *Wnt5*-*Vang* repulsive signaling.

We envision that Vang and Drl act cell autonomously to regulate axonal and dendritic guidance respectively and cell non-autonomously to modulate each other’s functions. Both Vang and Drl/Ryk have well-documented cell autonomous functions in neurite guidance. For example, vertebrate Vangl2 was localized to the filopodia of growth cones (35, 36) and *Drosophila* Vang mediates the repulsion of mushroom body axon branches in respond to Wnt5 (29, 37). Similarly, Drl and Ryk mediate the functions of Wnt5 and Wnt5a respectively in the targeting of dendrites (16, 38) and axons (5, 14, 17, 39, 40). Both proteins also have well-documented cell non-autonomous functions. Vertebrate Vangl2 acts as a ligand to steer migrating neurons (41, 42). We and others have shown that Drl could sequester Wnt5 using its extracellular Wnt Inhibitory Factor (WIF) motif (38, 43–45). Indeed, in this manner Drl may reduce Wnt5-Vang interaction, thus neutralizing Wnt5 repulsion of the glomeruli. How would Drl convert a glomerulus’s response to Wnt5 from repulsion to attraction? Increasing Ryk levels were proposed to titrate out Fz5 in chick retinal ganglion axons, thus converting growth cone response to Wnt3 from attraction to repulsion (46). Whether Drl function through a similar mechanism in the DA1 dendrites will require further investigation.

The opposing functions of Vang and Drl in a migrating glomerulus satisfies Geirer’s postulate for topographic mapping, which states that targeting neurites must detect two opposing forces in the target so that each neurite would come to rest at the point where the opposing forces cancel out (47). Thus, the relative levels of Drl and Vang activities in a glomerulus may determine its targeting position in the Wnt5 gradient (Fig 11). For the DA1/VA1d glomerular pair, the relative levels of Drl and Vang activities would result in the migration of DA1 up the gradient and VA1d down the gradient, that is, the rotation of the glomerular pair. The opposing effects of Wnt5 on the targeting glomeruli could allow the single Wnt5 gradient to refine the pattern of the olfactory map.

The rotation of the DA1/VA1d glomeruli bears intriguing resemblance to the PCP-directed rotation of multicellular structures such as mouse hair follicles and fly ommatidia, whose mechanisms remain incompletely understood (48–50). Our demonstration of the push-pull effect of Wnt5 on the glomeruli suggests that similar mechanisms may be involved in other PCP-directed rotations. Planar polarity signaling has emerged as an important mechanism in the morphogenesis of many tissues (20, 51, 52). However, apart from the molecules of the core PCP group (Vang, Prickle, Frizzled and Dishevelled) the identities of other signaling components are subjects of debate. A key question is the extracellular cue that aligns the core PCP proteins with the global tissue axes. Although Wnt ligands have been implicated, a definitive link between them and PCP signaling has been difficult to establish (20, 21, 53). Our work showing that Wnt5 and Vang act together to direct the orientation of nascent glomeruli adds to two other reports (54, 55) that Wnt proteins play instructive roles in PCP signaling. Another debate surrounds the role of Drl/Ryk role in PCP signaling. First identified as signal transducing receptors for a subset of Wnt ligands in *Drosophila* (17, 56), vertebrate Ryk was subsequently shown to act in PCP signaling (57, 58). These reports, combined with the lack of classical wing-hair PCP phenotypes in the *drl* mutant, led to the proposal that Ryk’s PCP function is a vertebrate innovation (59). Our demonstration of *drl*’s role in *Wnt5*-*Vang* signaling suggests, however, that the Drl/Ryk’s PCP function is likely to be evolutionarily ancient.

## Materials and Methods

### Transgenic animals and Crispr/Cas9 knockout of the *drl* gene

All transgenic fly lines were obtained from the Bloomington *Drosophila* Stock Center except for *UAS-Vang*, which was a gift from B. A. Hassan. To generate a new *drl* null allele on the *Mz19-Gal4* chromosome, exons 2, 3 and 4 of the *drl* locus (encompassing ∼90% of the *drl* open reading frame) were excised from the chromosome by Crispr/Cas9-mediated deletion using the sgRNAs GACAAGTGAAGGGGTGCTGT and GACACCTGTAGTGAGAGGTA following a published protocol (60). Ten individual offspring from Crispr/Cas9 fathers were crossed to *Adv/CyO* virgins to establish lines. The lines were screened by PCR using deletion-spanning primers to identify potential *drl* mutants. The PCR products were sequenced and one mutant, *drl^JS^*, with the expected precise deletion of the *drl* locus in the *Mz19-Gal4* background was chosen for study. The *drl^JS^* mutation failed to complement the AL phenotype of the *drl^2^* mutation, consistent with *drl^JS^* being a null allele.

### Clonal analyses

To induce *Vang* mutant ORNs, adults of the following genotype, *ey-FLP/+; FRT42 w^+^ cl/Mz19-Gal4 FRT42 Vang^6^ or (+)*, were obtained and dissected. To induce *Vang* and *drl* mutant PNs, the MARCM technique was employed (28). Third instar larvae of the following genotypes: *hs-FLP UAS-mCD8::GFP/+; FRT42 tub-Gal80/FRT42 GH146-Gal4 Vang^6^ or (+)* and *hs-FLP UAS-mCD8::GFP/+; FRT40 tub-Gal80/FRT40 Mz19-Gal4 drl^JS^ or (+); UAS-mCD8::GFP/+* were heat-shocked at 37°C for 40 minutes. Adult brains were dissected and processed as described below.

### Immunohistochemistry

Dissection, fixing and staining of adult or pupal brains were performed as previously described (27, 61). Rabbit anti-DRL (1:1000) was a generous gift from J. M. Dura; rat anti-Vang (1:500) was a gift from D. Strutt; mAb nc82 (1:20; (62) was obtained from the Iowa Antibody Bank; rat anti-mCD8 mAb (1:100) was obtained from Caltag,. The secondary antibodies, FITC-conjugated goat anti-rabbit, Cy3-conjugated goat anti-mouse and FITC-conjugated goat anti-rat, were obtained from Jackson Laboratories and used at 1:100 dilutions. The stained brains were imaged using a Zeiss 710 confocal microscope

### Quantification of glomerular rotation

Two different quantification strategies were employed. To quantify the displacements of single glomeruli labeled by the *Or-Gal4* drivers (6), the angle subtended at the VA6 glomerulus (close to the center of the AL) by the dorsal pole and the labeled glomerulus (in the dorsal → lateral → ventral → medial direction) was measured. To quantify the rotation of the DA1 and VA1d glomeruli (22) around each other, the angle subtended at the VA1d glomerulus by the dorsal pole and the DA1 glomerulus (in the dorsal → lateral → ventral → medial direction) was measured. Data were collected, analyzed and plotted using the Prism statistical software. For two-sample comparisons, unpaired Student’s *t*-tests were applied. For comparisons among more than two groups, one-way AVOVA tests were used followed by Tukey’s test. Rose diagrams were plotted using the Excel program.

## Acknowledgments

We thank L. Luo and the Bloomington Stock Center for providing fly stocks, J. M. Dura and D. Strutt for their generous gifts of the anti-Drl and anti-Vang antibodies respectively, and J. N. Noordermeer and J. M. Dura for their comments on the paper. This work was supported by grants from the NIH/NIDCD (DC010916-02A1) awarded to H. Hing.

## Author Contributions

N.R. identified *Vang* from a screen for mutations that disrupted AL development. L.G.F. constructed the *pCFD4-drl^KO^ Crispr* transgene. J.S. carried out the Crispr/Cas9 genetic crosses to create the *drl^JS^* allele and characterized it molecularly. H.H. carried out all the experiments characterizing *Vang* and *drl* as well as establishing *Vang* as a component by which *wnt5-drl* signaling orients the rotation of the glomeruli. H.H. analyzed the data; and H.H. and L.G.F. wrote the manuscript with contributions from the other authors.

